# Bayesian learning ecosystem dynamics with delayed dependencies from incomplete multiple source data : an application to plant epidemiology

**DOI:** 10.1101/453050

**Authors:** Stéphane Dupas

## Abstract

Ecosystem dynamics forecasting is central to major problems in ecology, society, and economy. The existing models serve as decision tools but their parameters valitity are usually not confronted to real data in a formalized approach. Dynamics bayesian network inference is promissing but limited when dealing with incomplete multiple source time series with delayed time dependencies. We propose here a temporal bayesian network with time delay and aproximate inference algorithm, to learn altogether cryptic ecosystem variables, missing data, and model parameters. The novelty in the approach is that it combines simulation-based and likelihood-based aproximate bayesian inference. The advantage of simulation based is that it allows to sample hidden processes. The advantage of likelihood based is that it provides a summary statistics that is really representing the model we are interested in. The ecosystem variables and the missing data are simulated from indicator variables using the probabilistic indicator-ecosystem model. The likelihood is estimated by averaging the probability of observed-simulated data over simulations, the parameter space is sampled with Metropolis Hasting algorithm. Another innovative proposition is to parametrize the network structure in order to learn model structure within a space provided by prior distribution. We apply to plant epidemiology.

## Introduction

The systems we live in, be they natural or human constructions, are ecosystems. Understanding and predicting their dynamics is central to many of our stakes on earth. We can model their emerging properties from the knowledge of the interactions, but this pose problems of extrapolation and it is more difficult to analyse interactions separately than to study the system as a whole and infer interactions from node dynamics. Yet, this requires methods for statistical inference. Many studies attempts to infer interaction from statistical relationship in static data, but the causal inference can only be attained from temporal relationships among system nodes.

For deterministic linear models such as lotka voltera, systems parameters solutions can be obtained by solving systems of equations through matrix algebra [1].

When dealing with statistical inference of probabilistic ecossytem dynamics statistical inference, the most well polished techniques are Kalman filters for continuous state linear dynamic systems (LDS) and Hidden Markov Models (HMMs) for discrete state sequences. But real world ecosystems are determined by the interaction between variables of different nature (information, energy and material) that can be discrete or continuous. In such hybrid bayesian network, the main solution is to discretize continuous variable and use discrete model inference technique. Truncated exponential has been proposed to better approximate continuous variable discretization [2]. Note that inference of DBN structure from discrete or discretized data is promising but has to be interpreted with caution [3]. Another front of research is nonlinear relationships between variables. It has been approximated as a succession of linear models indexed by switching variable (Switching linear dynamic systems, SDLS) [4]. Particle filter monte carlo methods acompanied with resampling have been proposed to treat, in principle any type of distribution, non linearity and non stationarity [5, 6]. When possible, Rao blackwellized marginalization can treat problems of sampling in high dimensional space [7]. These particle filters and sequential Monte Carlo methods can allow to infer posterior bayesian distribution

Another issue is the time delay dependance. In DBN, each time slice depend only on the previous time slice variables (*X_t_* depends only on *X_t_*−1). However time delays are comonly oberved in ecosystem dynamics and stability properties are highly dependent on time lag and turnover rates properties of the system [8]. Time lags are essential to predict important properties of ecosystems, in particular collapsing risk. A framework for learning temporal bayesian network with deeper time dependencies would be useful.

Finally most of the data set for ecosystem dynamics inference contain missing data time series data is usually incomplete. And the probability that some of the measures in the time series is lacking increases exponentially with the number of data. Missing data can include a whole variable. Variables interacting ecosystems and constituting the ecosystem can be unmeasured but estimated though indicators or samples.

When considering a new ecosystem, a specific mathematical and software solution may or may not be found. A generic inference tool allowing to encompass the different ecosystem types would however be of interest.

Aproximate Bayesian Computation is now comonly used for model inference in ecology and evolution [9]. Parameters are sampled from prior distribution, data are simulated and summarized with statistics and the parameters that gave the summary statistics closest to the data are conserved in the posterior [10, 11]. It has been suggested of use for modelling dynamic bayesian networks. In many ecosystem dynamics, simulating time series based on model parameters would bring a lot of noise in summary statistics and reduce method power due to historical contigency. Like for kahlman filter it would be interesting to update historical simulations with data in real time. Another issue with ABC is the choice of summary statitstics. In this work we propose to combine likelihood and Monte Carlo data simulation to get the advantage of each. Ecosystem variable needed for likelihood calculation are simulated from observation and model altogether. Then posterior model parameters can be sampled using likelihood averages times prior probability.

Example of plant epidemiological models. Use prior information of known epidemiological model [12]

## Probabilistic model

Let us consider the data 
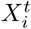
 representing several incomplete times series where *X_i_*is the variable, and *t* the time when it was sampled. 
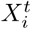
 are directly related to underlying interacting ecosystem variables 
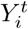
. *X* represent the measured variables and *Y* represent the interacting variables in the ecosystem. For instance, 
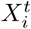
 is an observation of a species at time *t* and 
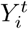
 is the density of this species at time *t*, or, 
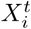
 is an event of pesticide application in a field, and 
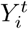
 is the concentration of pesticide residues in the leaf at time *t*. We assume we can define a recursive form of the ecosystem model which parameter are time independent. Note that to model the exosystem dynamics among *Y_i_* we may need additional hidden variables *Z_j_*. For instance the fungus targetted by the pesticide can have a criptic density 
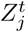
 at latent development stage within leaves cells where it cannot be observed directly [12]. In addition, for the observed variables 
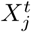
 there can be periods with missing data. The difficulty in learning such dynamics models is to learn hidden variables *Z* and unobserved variables at all time. In order to get rid of these latent variables we propose a recursive sheme with deeper time dependencies than from *t* to *t* + 1 so that we focus on modeling dependencies among variables directly linked to observations. The rationale of our approach is to be able to adapt machine learning to any kind of data and ecosystem.

The solution I propose in this paper to this problem of inference of non linear probabilistic ecosystem dynamic model with hidden variable and deep time dependence from incomplete data is to combine data simulation with likelihood. First the ecosystem variables are simulated from indicator data using indicator and ecosystem models, then joints probability of observed and simulated data is calculated and aeraged over simulations for each prior sample. The algorithm is not going to learn hidden variables but to simulate them. Data completed by simulations will allow to calculate an estimate of the likelihood of the model. If the residual variation of the simulated data is not too important averaging over an acceptable number simulations can provide estimates of model likelihood and permit bayesian inference without to make use of non heuristic summary statistics as in Aproximate Bayesian Computation. I name this approach, MCL (Monte Carlo likelihood), or bayesian computation with monte carlo likelihood.

Let us consider the ecosystem dynamics model presented in figure 1. It is consituted with a time series of indicators 
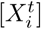
 collected at time *t* for variable *X_i_* related to ecosystem variables 
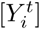
. *p*(*X/Y, θ_X,prior_*) is the sampling distribution model of the indicator variables, and *p*(*Y*/Γ(*Y*), *θ_Y,prior_*) is ecosystem dynamics model of underlying ecosystem variables described in figure 1 for its unrolled version, where Γ(*Y*) are the parents of *Y*. Note that Γ(*Y*) can be given as prior or parametrized through *θ_prior_* The model *θ,* Γ(*Y*) is time independant but not markovian since *Y_t_* does not depend only on *Y_t−1_*. Time dependence can also be delayed with Γ(*Y*) given by *θ* value. In the case of potato late blight disease the fungus development includes a latent period where development is cryptic in the leaves and it’s survival is not susceptible to environmental conditions (Figure 2). Sporangium produced and infesting leaves the day before the sampling will not be observable as mycelium on the leaves. Since the time dependance can be delayed, these cryptic variables can be analytically removed from the model with time delayed responses. In the case of potato late blight disease, the cryptic variable is the latent stage of the fungus, that give birth to exposed micelium outside the leave and originated from infestation by sporangium a few days before (the development time depends on the temperature). We can propose a prior function relating the number of newly emerged mycelium from past infesting sporangium as a result of the number of sporangium at a period before which length depend on development rate of the latent stage, and its infestation rate at that stage, that depends on climatic conditions. We recomend removing analytically any criptic variable from the model.

**Figure 1.**
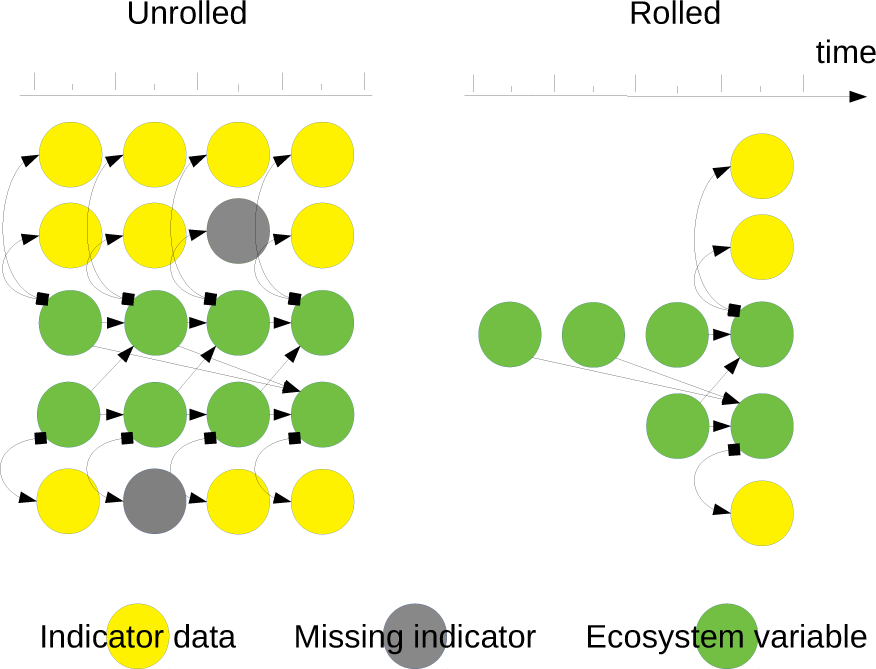
Generic ecosystem - indicador model

**Figure 2.**
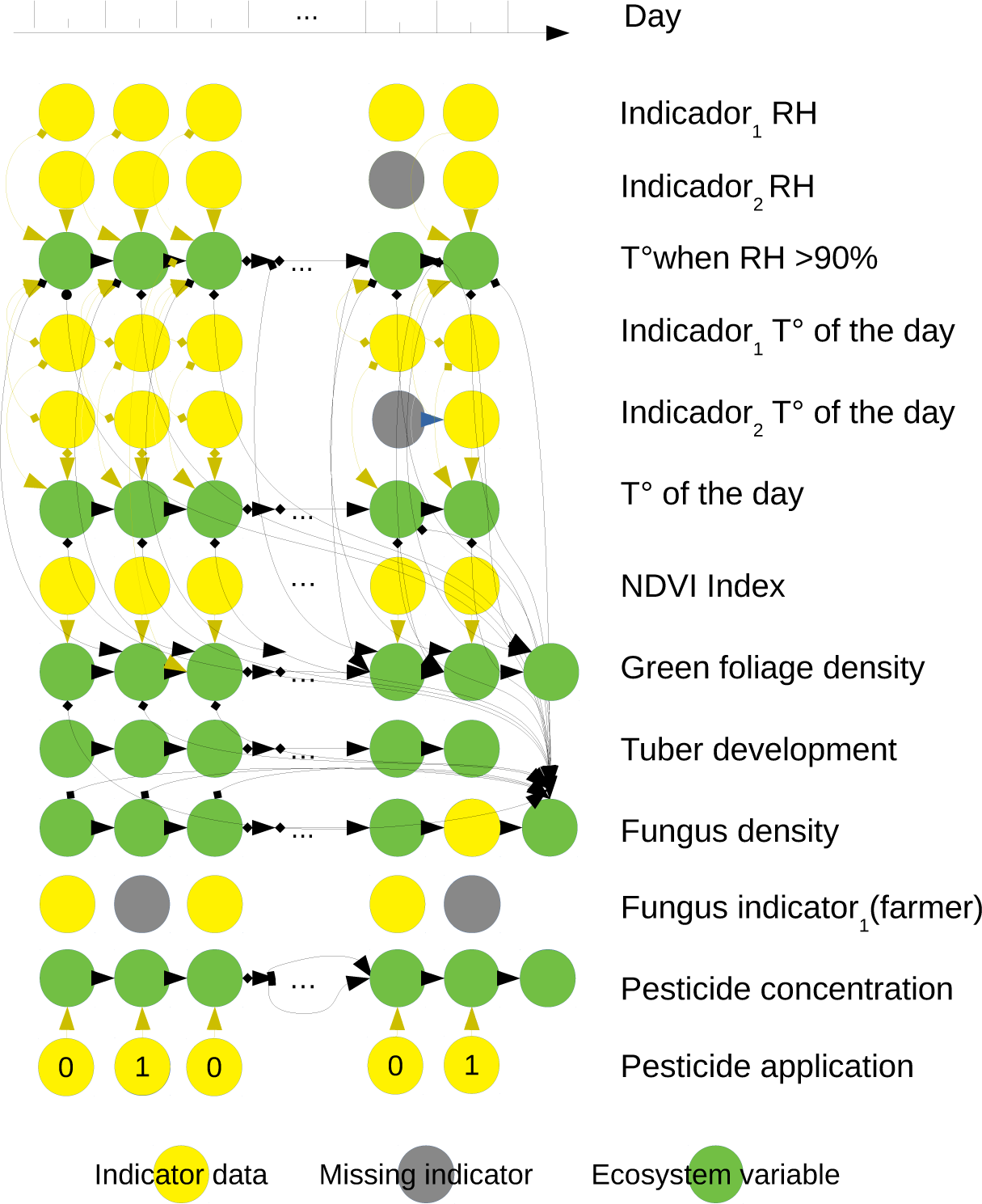
Potato late blight ecosystem model. We distinguish between indicator variables in yellow and ecosystem variables in green. Indicator variable and ecosystem variable are linked by sampling or indicator probabilistic model through bayesian rules (yellow arrows). Ecosystem variable are linked together by ecosystem temporal bayesian network model with deep time dependence (black arrows). Nodes in grey are undelying missing data

We assume we can define a prior model *p*(*Y/θ_Y_,* Γ(*Y*)) for ecosystem variable dynamical interactions with prior probability distributions for *θ_Y_*. We assume we can define a prior model for measured variable sampling 
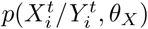
 where *θ_X_* are sampling parameters. For instance for temperature they determine the precision of the thermometer or the difference between the temperature at the plant leaves its value in the meteorological station. The model can also include deterministic fixed parameters.

The likelihood for a sample prior will be calculated by averaging probability of simulating ecosystem data from indicator and ecological sample prior as detailed in Dupas et al unpublished.

Condider *x* indicator data. If we simulate *y* from *p*(*y/x, θ_X_, θ_Y_*), then model likelihood is (Dupas et al unpublished)

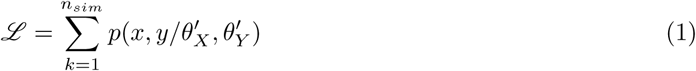

### Algorithm

1. Discretize spatio-temporally the data
2. Define indicator prior sampling model *p*(*X/Y, θ_X_*) and ecological prior sampling model structure Γ(*Y*) and probability distribution *p*(*Y*/Γ(*Y*),*θ_Y_*)
3. Define parameter proposal rules of metropolis hasting sampler
4. Sample 
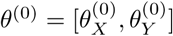
 from their prior
5. Repeat for *j* ∈ [1‥*n_generations_*] :

a. define *θ^g′^* from *θ*^(*j*)^using proposal rules
b. if *j* > 0 set *p* = *p^′^*
c. Repeat for *s* ∈ [1‥*n_simulations_*] :

i. For *t* ∈ [*t_min_, t_max_*] and for *i* ∈ [1‥*n_variables_*], complete time series data by simulating 
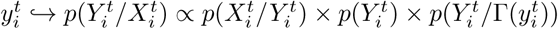
, where 
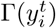
 represent the parents of 
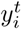
, already simulated.
ii. Calulate (*x, y*) probability 
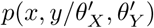
 over the completed time series
d. estimate the likelihood as the average of (*x, y*) probability among simulations 
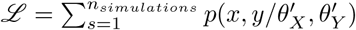
e. Calculate posterior 
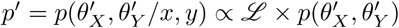
f. if *j* > 0 Calculate 
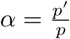
 and

i. if *α* ⩾ 1 set *θ*^(*i*)^ = *θ^′^*
ii. if *α* < 1 set *θ*^(*i*)^ = *θ^′^* with probability *α*
iii. if *thining* | *j* then save *θ* in *θ_sample_*
6. return *θ_sample_*

## Plant epidemiology example : the potato late blight

I will propose non linear probabilistic models for the different components of the likelihood in the case of potato late blight disease (figure 2). Plant epidemiology depends on historical and environemental constraints which are very specific to each locality. We need to be able to learn from these specificities in real time.

In this model we consider the prior information of *Phythophthora infestans* life cycle relationships with environmental variables proposed by [12] where the emidemics is determined by

1. *rLGR*, the relative lesion growth rate, function of *T*, the mean daily air temperature
2. *rSR*, the sporulation rate function of *T H*, the mean daily air temperature when relative humidity is above 90%,
3. *LP*, the latent period, or the period of intracellular development during which the disease is undetectable in the leaf tissues, function of *T*.

In [12] the relationship between these parameters and climatic variables are polynomial of second or third order with 3 or 4 parameters each.

## Acknowledgments

This work has been supported by IRD EGCE research unit project and BIO INCA international mixed laboratory

